## Supplementary figures and images for "Rab11a mediates cell-cell spread and reassortment of influenza A virus genomes via tunneling nanotubes"

### Supplemental Figure 1A

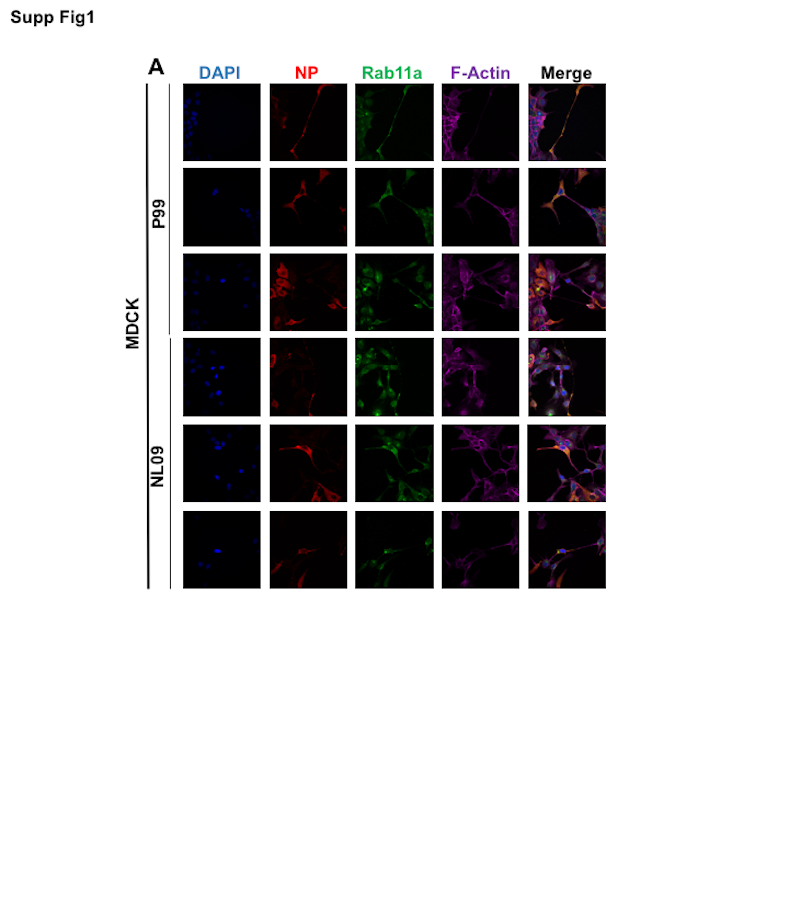

### Supplemental Figure 1B

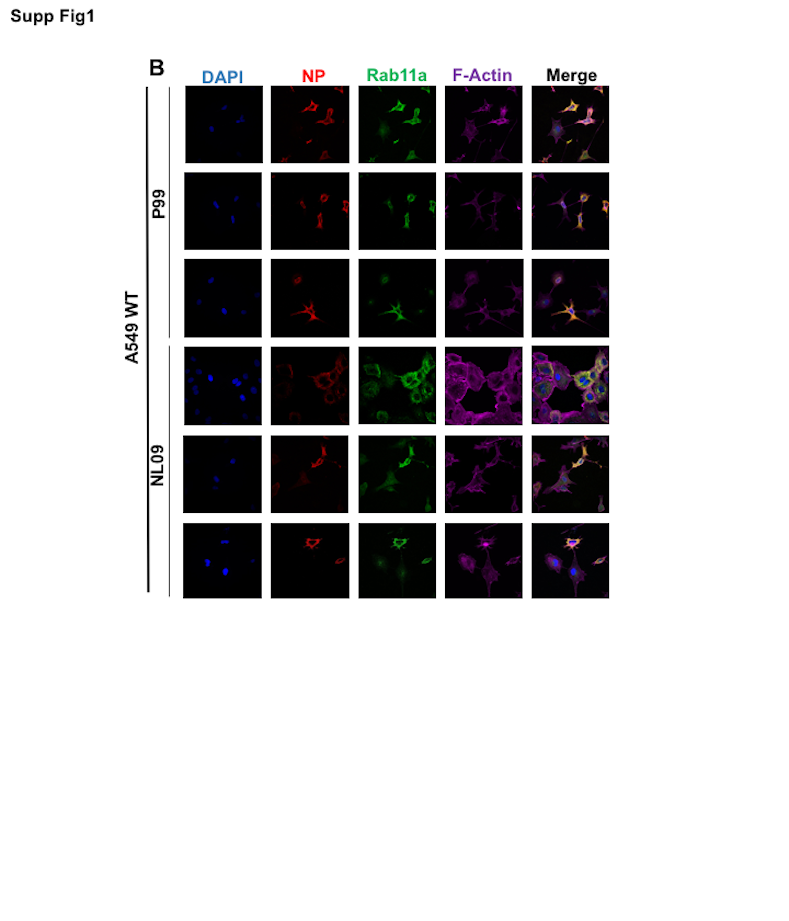

### Supplemental Figure 1C

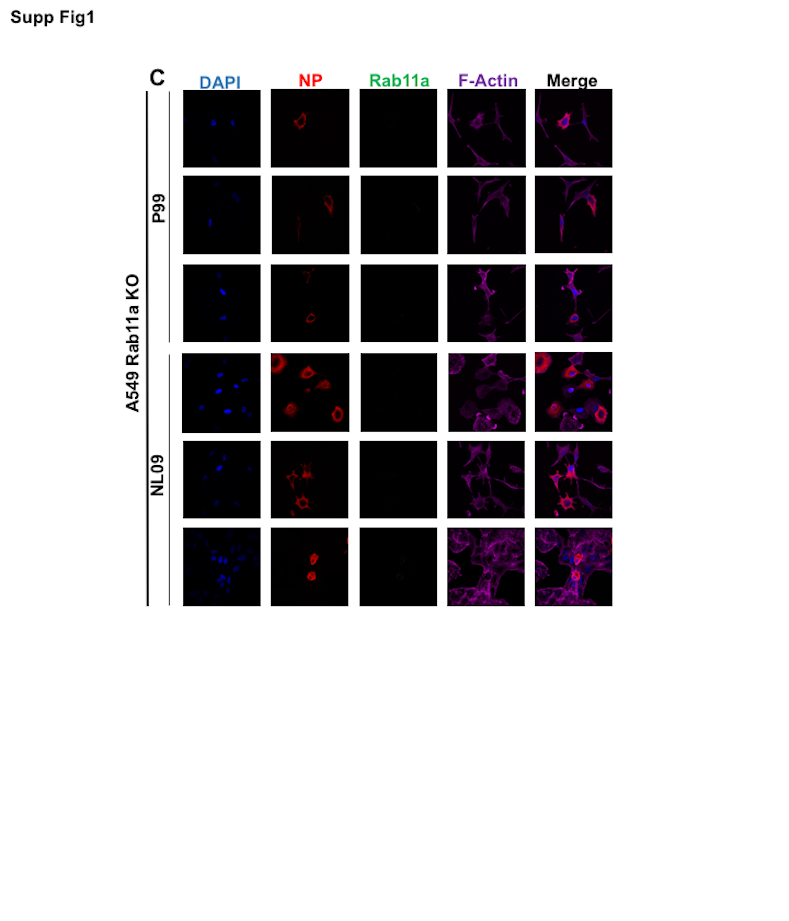
